# Anosognosia for Hemiplegia as a tripartite disconnection syndrome

**DOI:** 10.1101/560326

**Authors:** V. Pacella, C. Foulon, P.M. Jenkinson, M. Scandola, S. Bertagnoli, R. Avesani, A. Fotopoulou, V. Moro, M. Thiebaut De Schotten

**Author notes:** Equal contribution (A. Fotopoulou, V. Moro, M. Thiebaut De Schotten). Corresponding Author &.

## Abstract

The rare syndrome of Anosognosia for Hemiplegia (AHP) can provide unique insights into the neurocognitive processes of motor awareness. Yet, prior studies have only explored predominately discreet lesions. Using advanced structural neuroimaging methods in 174 patients with a right-hemisphere stroke, we were able to identify three neural networks that contribute to AHP, when disconnected: the (1) premotor loop (2) limbic system, and (3) ventral attention network. Our results suggest that human motor awareness is contingent on the joint contribution of these three systems.

---

Motor awareness allows individuals to have insight into their motor performance, a fundamental aspect of self-awareness. However, following damage to the right hemisphere, patients with left paralysis may show delusions of intact motor ability, or anosognosia for hemiplegia (AHP, 1). Hence, studying AHP offers unique opportunities to explore the neurocognitive mechanisms of motor awareness.

While early studies regarded AHP as secondary to concomitant spatial deficits such as hemineglect 2 caused by parietal lesions, more recent experimental and voxel-based, lesion-symptom mapping (VLSM) results suggest that AHP is an independent syndrome. These earlier studies address AHP as an impairment of action and body monitoring, with lesions to the lateral premotor cortex and the anterior insula (3,4), affecting patients’ ability to detect discrepancies between feed-forward motor predictions and sensorimotor feedback. However, these hypotheses are insufficient to explain all the AHP symptoms, such as patients’ inability to update their beliefs based on social feedback or more general difficulties experienced in their daily living (5,6). Indeed, others have suggested that AHP can be caused by a functional disconnection between regions processing top-down beliefs about the self and those processing bottom-up errors regarding the current state of the body (5,7). Nevertheless, to date the brain disconnection hypothesis could not be explored due to the relatively small sample size and the standard methodology of previous studies, which favours the implication of discreet lesion locations in the pathogenesis of AHP.

Here, to overcome this gap, we took advantage of (1) the largest cohort of AHP patients to date (N = 174; 95 AHP patients diagnosed by (8) and 79 hemiplegic controls) and (2) an advanced lesion analysis method (BCBtoolkit, 9). This method generates a probabilistic map of disconnections from each patient’s brain lesion to identify the disconnections that are associated with given neuropsychological deficits at the group level. Previous use of this connectivity approach has already proven fruitful in the study of neuropsychological deficits (10–12).

We predicted that AHP would be associated not only with focal grey matter lesions, but also with long-range disconnections due to the white matter damage, in particular to tracts associated with sensorimotor monitoring and self-reflection. Specifically, we anticipated the possibility that motor awareness emerges from the integrated activation of separated networks (13,14), whose contributions feed into the multifaceted expression of the syndrome.

## Results

To test these predictions, we first conducted anatomical investigations to identify lesion sites and created probability maps of white matter tracts' disconnection. These results were statistically analysed by means of regression analyses, to identify the contribution of grey and white matter structures in AHP, taking into account differences in age, lesion size, lesion onset-assessment interval and critical motor and neuropsychological deficits (i.e. covariates of non-interest). Considering our sample size and a power of 95%, t values above 2 correspond to a medium effect size (cohen d > 0.5) and t values above 3.6 correspond to a large effect size (cohen d > 0.8).

The regression computed on the lesion sites (**Figure 1a**) indicated the involvement of grey matter structures previously associated with AHP (15), such as the insula (anterior long gyrus, t = 4.89; p = 0.002), the temporal pole (t = 4.77; p = 0.003), and the striatum (t = 4.68; p = 0.003) as well as a very large involvement of white matter (t = 4.98; p = 0.002). The second regression on white matter maps of disconnection (**Figure 1b**) revealed a significant contribution of the cingulum (t = 3.85; p = 0.008), the third branch of the superior longitudinal fasciculus (SLF III; t = 4.30; p = 0.003), and connections to the pre-supplementary motor area (preSMA; t = 3.37; p = 0.013), such as the frontal aslant and the fronto-striatal connections.

**Figure 1:**
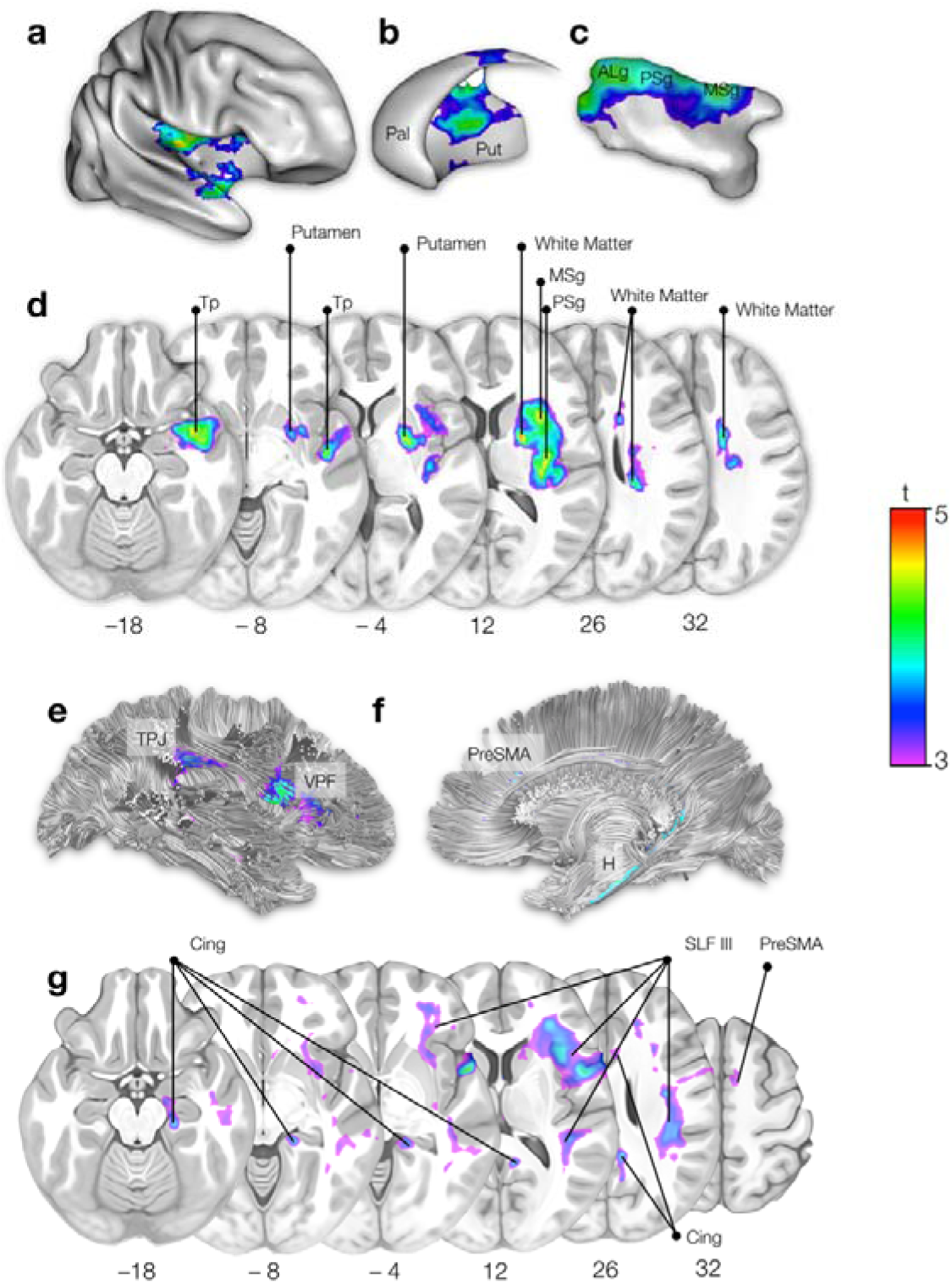
On the top half, statistical mapping of the lesioned areas in AHP. a) right hemisphere b) striatum c) insula d) axial sections. Pal: pallidum; put: putamen; ALg: anterior long gyrus; PSg: posterior short gyrus; MSg: middle short gyrus; Tp: temporal pole. On the bottom half, statistical mapping of the brain disconnections in AHP. e) right hemisphere lateral view; f) right hemisphere medial view; g) axial sections. TPJ: temporo-parietal junction; VPF: ventral prefrontal cortex; preSMA: pre-supplementary area; H: hippocampus; Cing: cingulum; SLF III: third (ventral) branch of the superior longitudinal fasciculus; PreSMA: pre-supplementary motor area.

To test whether AHP emerges from the disconnection of each of these networks independently or together as a whole, we investigated the contribution pattern of the tracts’ disconnection to AHP (individual or integrated), by means of Bayesian computation of generalised linear multilevel models. All of the possible binomial models (n=95) were computed, starting from the null model (with only the covariates of non-interest) to the full model, with all the covariates of non-interest, the tracts, and all the interactions among them (16) (see Methods section). The results confirmed that the disconnection of each tract is critical to AHP (Cingulum, BF_10_ = 270.98; FST, BF_10_ = 180.48; FAT, BF_10_ = 367.61; SLF III, BF_10_ = 571.49). However, results indicate that the model that best fits with our data (99% of probability) includes the contribution of all the four tracts (**Figure 2**).

**Figure 2:**
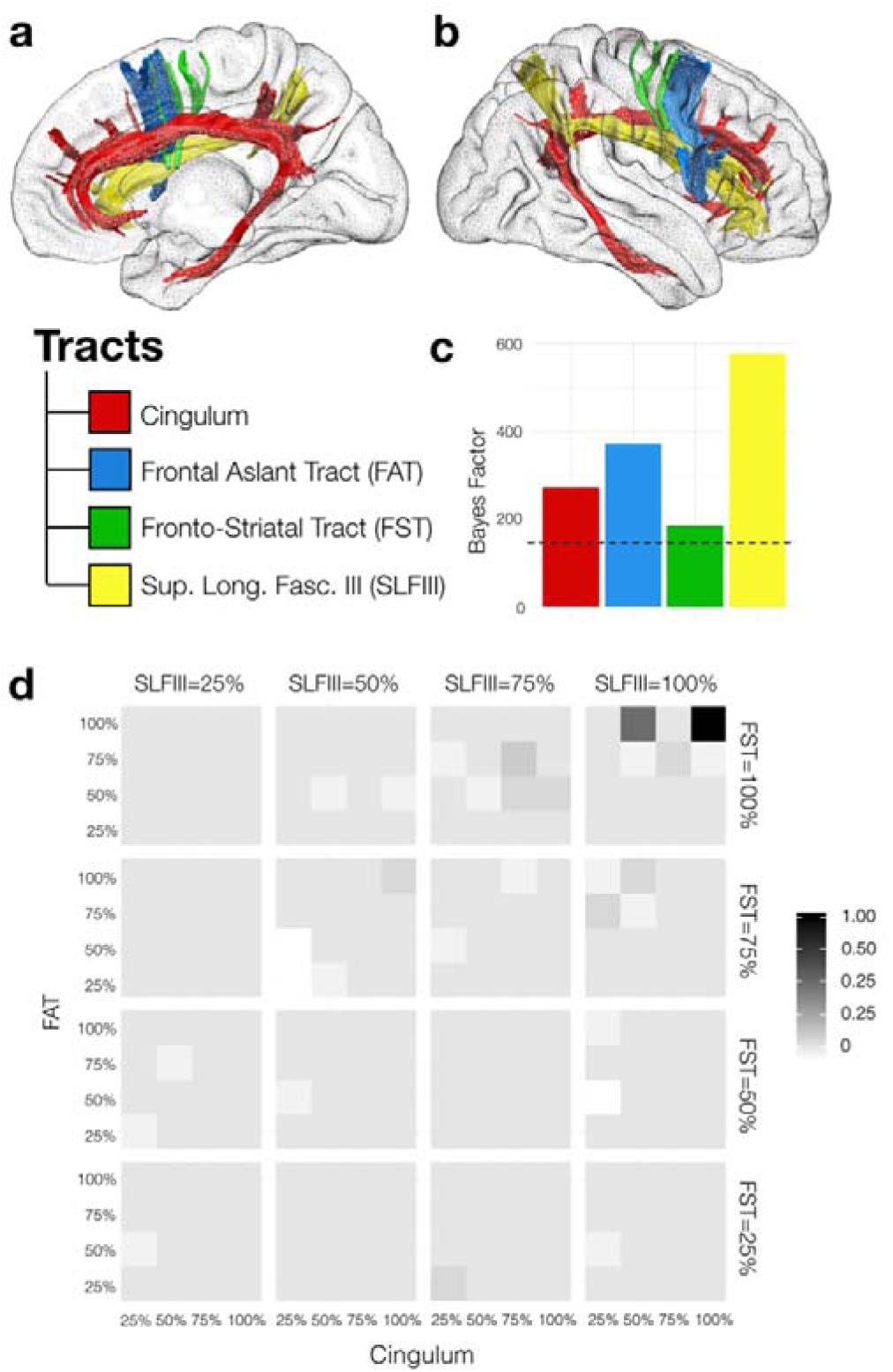
Motor awareness network a) right hemisphere medial view; b) right hemisphere lateral view c) Bayes Factors for the four models, each one representing the hypothesis that the presence of disconnection in a tract is necessary to explain AHP, against the null model (i.e. no disconnection is necessary to explain the presence of AHP). The dashed line represents BF_10_ = 150, the boundary suggested by (17) as the decision criteria for very strong support of hypotheses. The plot shows that each tract contributes to AHP symptoms. d) proportion of AHP over non-AHP patients, normalised to its maximum (20). Darker the line, the greater the presence of AHP compared to non-AHP. The disconnections of tracts are divided into four categories: 25% - (0, 25]; 50%: (25, 50]; 75% - (50, 75] and 100% - (75, 100]. In each subplot, the y-axis shows the percentage of FAT disconnection and the x-axis shows the percentage of Cingulum disconnection. The subplots are divided horizontally by the percentage of SLF III disconnection, and vertically by the percentage of FST disconnection. In the all squares, binomial tests were computed to check if the number between AHP and non-AHP is different. The only significant difference (p=.01, the number of AHP is greater) is when the probability of disconnection for each tract is: SLFIII = 100%, FST = 100%, Cingulum = 100% and FAT = 100% (the black square on the extreme top-right).

These results, derived from the largest lesion mapping study on AHP to date, show that white matter disconnections in three networks contribute to AHP: (1) posterior parts of the limbic network (i.e. connections between the amygdala, the hippocampus and the cingulate gyrus); (2) the ventral attentional network (i.e. connections between temporo-parietal junction and ventral frontal cortex), through the SLF III; and (3) the premotor loop (i.e. connections between the striatum, the preSMA and the inferior frontal gyrus).

## Discussion

Previous lesion mapping studies in AHP have highlighted the role of discrete cortical lesions in areas such as the lateral premotor cortex or the insula, and suggested corresponding theories of motor and body awareness (3,4 and 5,6, for a critical review). By contrast, our results suggest that AHP is a tripartite disconnection syndrome involving disruptions in networks that include, but also extend beyond, sensorimotor circuits. Correspondingly, motor awareness should be regarded as the collaborative effort (i.e. integration of a number of cognitive processes, rather than a purely motor monitoring function. Indeed, this interpretation is consistent with the delusional features of AHP (see 18) and a variety of experimental findings in AHP, such as the fact that awareness can be influenced by mood (19,20), or perspective-taking (21,22).

Specifically, the cingulum connects limbic system structures that have been previously associated with emotional and memory processing, and is part of the default mode network (23)–a pattern of intrinsic connectivity observed during self-referential, introspective states, including autobiographical retrieval, future imaging and mentalisation. These abilities relate to well-documented deficits in AHP patients’ general awareness (“why are you in hospital?”), anticipatory awareness (“are you able to reach the table with your left hand?”, 24) and mentalisation (25; “the doctors think there is some paralysis, do you agree?”, 26).

The ventral attentional network (i.e. SLF III connections between temporo-parietal junction and ventral frontal cortex, as well as lesions in the insula and temporal pole) reorients attention towards relevant stimuli (27). This disconnection prevents the possibility to appreciate the bottom-up stimuli referring to one’s own paralysis and (along with the limbic system) to update beliefs regarding the current body’s condition (“I can walk as I have always done”; “I have just clapped my hands”, 21). The insula is also crucial in these processes due to its important role in integrating external sensory information with internal emotional and bodily state signals (28).

Finally, the observed disconnections of the pre-motor network (pre-SMA, striatum and inferior frontal gyrus) suggest difficulties in monitoring motor signals and learning from action failures (“did you execute the action? Yes, I have”).

Crucially, none of these networks (i.e. the limbic system, the ventral attentional network, and the premotor system) alone can fully explain AHP (**Figure 2c**). It thus appears that bottom up deficits in interoceptive and motor salience monitoring need to be combined with higher-order deficits, collectively leading to a multifaceted syndrome in which premorbid beliefs and emotions about the non-paralysed self dominates current cognition about the paralysed body.

These results open up interesting hypotheses on the hierarchical or parallel relations between the three networks, in terms of temporal activations (either serial, parallel or recurrent) that remain to be explored in future studies.

The main limitation of the study is related to manual lesion delineation and registration methods (29,30) and the sensitivity level of neuroimaging techniques that do not depict the full extent of damage produced by stroke lesions (31). However, these limitations mainly apply to small sample studies, while here, the large number of patients investigated reduces these risks.

In conclusion, on the basis of a large (N = 174) and advanced lesion-mapping study, we demonstrate a tripartite contribution of disconnections to the pre-motor network, the limbic system, and the ventral attentional network to motor unawareness. We thus suggest that motor awareness is not limited to sensorimotor attention and monitoring but also requires the joint contribution of higher-order cognitive components.

## Acknowledgements

We thank Lauren Sakuma for useful discussion and edits to the manuscript. This project has received funding from the European Research Council (ERC) under the European Union’s Horizon 2020 research and innovation programme (grant agreement No. 818521). This study was supported by a European Research Council Starting Investigator Award (ERC-2012-STG GA313755, to A.F.); Fondation pour la Recherche Médicale (FRM DEQ20150331725); by MIUR Italy (PRIN 20159CZFJK), and University of Verona (Bando di Ateneo per la Ricerca di Base 2015 project MOTOS) (to V.M.).

## Materials and Methods

### 1. Design and Statistical Analysis

The aim of the study was to explore the white matter disconnections involved in AHP. To this end, we investigated the neural systems that contribute to the symptoms of AHP. To the best of our knowledge, this approach has never been applied to the study of AHP and it can shed light on the theoretical and phenomenological complexity of the disease, by integrating and going beyond existing findings gained through classic lesion studies (3,4,15,18,32,33).

For this purpose, we collected neuroimaging and clinical data from a large sample of right hemisphere stroke patients. To compute lesion sites and map of disconnections that were strictly related to the AHP pathology, we compared our target group of AHP patients with a group of stroke patients with hemiplegia but without AHP. Patients’ map of lesions and disconnections were statistically compared between the two groups and adjusted for covariates of non-interest. Variance related to patients’ demographic variables (age and education level) was removed. As previous lesion studies (15,32) found some differences in neuronal correlates of AHP in acute and chronic stages, the interval between lesion onset and neuropsychological assessment was considered as a covariate of non-interest as well. We also included the lesions’ size of our sample, taking into account the number of voxels of each lesion as a nuisance variable.

Finally, when computing the AHP map of lesions and disconnections we controlled for the clinical (onset-assessment interval, motor deficits) and neuropsychological (personal, external neglect and memory impairments) symptoms that are often associated with AHP but are related to different patterns of disconnection.

The tracts emerging from this analysis were further analysed by means of Bayesian models to confirm the individual involvement of each tract and test their joint contribution to AHP (see details below).

### 2. Patients

Data from 195 stroke patients with unilateral right hemisphere damage were collected from two collaborating centers based in Italy and the United Kingdom over a period of 10 years.

Patients’ inclusion criteria were: (i) unilateral right hemisphere damage, secondary to a first-ever stroke, as confirmed by clinical neuroimaging; (ii) severe plegia of their contralateral upper limb (AHP left arm, MRC ≤ 2), as clinically assessed (MRC scale). Exclusion criteria were: (i) previous history of neurological or psychiatric illness; (ii) medication with severe cognitive or mood side-effects; (iii) severe language, general cognitive impairment, or mood disturbance that precluded completion of the study assessments.

The MRI or CT neuroimaging data were available for 174 out of 195 patients. They were divided into two groups according to the presence/absence of AHP (see below for AHP assessment details), resulting in a group of 95 AHP patients and 79 non-AHP, hemiplegic control (HP) subjects. Among these, clinical and anatomical data of 40 AHP patients and 27 controls has been described in a previous study (15). Groups were balanced for demographic data (age, education, interval period between lesion and assessments) and lesion size. As the data were collected from different stroke recovery units, we took into account the neurological and neuropsychological tests that were most commonly administrated to all the patients across the different centers (**Table 1**).

**Table 1.**
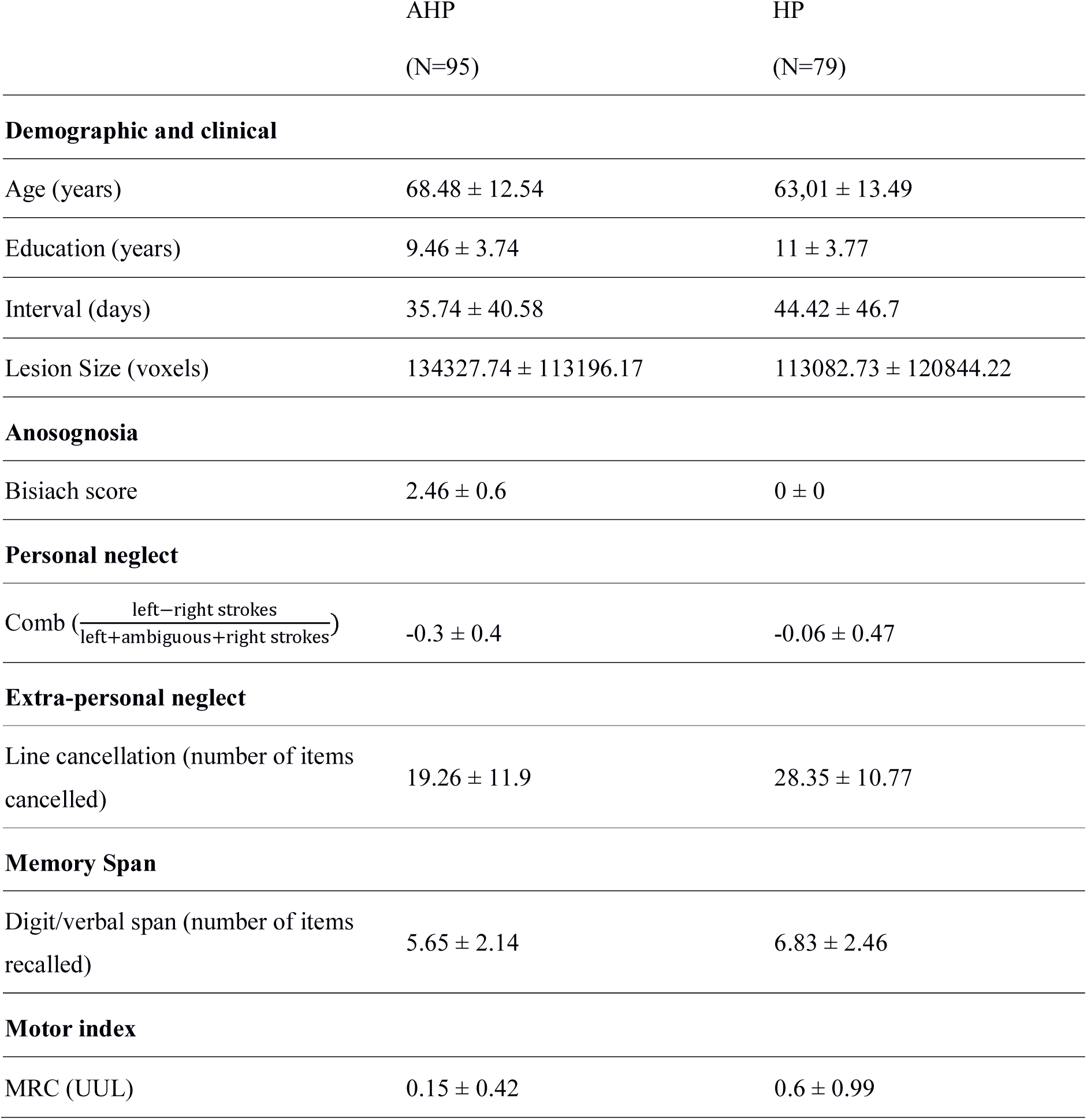
For AHP and control groups, mean and (± standard deviation) of demographic and clinical variables, neurological and neuropsychological assessments are reported.

All patients gave written, informed consent and the research was conducted in accordance with the guidelines of the Declaration of Helsinki (2013) and approved by the Local Ethical Committees of each center.

### 3. Neurological and Neuropsychological Assessment

Patients were identified as anosognosic or control according to their score in the Bisiach scale (8). This investigates the explicit form of awareness related to one’s limb paralysis. During the scale administration, patients were required to verbally answer a 4-point interview about their current condition: a ‘0’ score indicates a spared consciousness of the disease (= the disorder is spontaneously reported or mentioned by the patient following a general question about his/her complains), a ‘1’ score is assigned when patients refer to their disability only after specific questions about the strength of their left limbs, while patients scoring ‘2’ or ‘3’ are considered anosognosic for their awareness of the disease emerging only after a demonstration through a routine technique of neurological examination (score 2) or not emerging at all (score 3).

Plegia of the contralesional upper limb was assessed through the Medical Research Council 5-point scale (34), ranging from 5 (normal functioning) to 0 (no movement).

Personal neglect was assessed by means of the ‘Comb’ test, from the ‘Comb/Razor test’ (35). We referred to each patient’s score on the line cancellation subtest of the BIT as our measure of extra-personal neglect (Behavioral Inattention Test, 36). Finally, we used the digit/word span (37,38) to assess working memory. The 3-nearest neighbour computation replaced the missing data from the demographic and clinical variables (education: 4.9%; lesion-assessment interval: 0.05%; motricity index: 1.9%; personal neglect: 1.7%; extra-personal neglect: 2.5%; memory span: 6.2%).

In order to compare results expressed in different scoring ranges, all the scores from neuropsychological tests were transformed to z-scores, with higher scores corresponding to better performances.

### 4. Lesions drawing

Patients’ neuroimaging data was acquired via Computerized Tomography (CT) and Magnetic Resonance (MRI) and lesions were segmented and co-registered using the manual procedure already described by Moro and colleagues (15).

The lesion drawing was performed blindly and independently by two of the authors (VM, SB), prior (blind) to the group classification. In cases of disagreement on a lesion drawing, a third anatomist’s opinion was consulted (<10%).

Scans were registered on the ICBM152 template of the Montreal Neurological Institute, furnished with the MRIcron software (ch2, http://www.mccauslandcenter.sc.edu/mricro/mricron/). First the standard template was rotated on the three plans (size: 181 x 217 x 181 mm, voxel resolution: 1 mm^2^) in order to match the orientation of patient’s MRI or CT scan. Lesions were outlined on the axial slices of the rotated template. The resulting lesion volumes were then rotated back into the canonical orientation, as to align the lesion volumes of each patient to the same stereotaxic space. Finally, in order to remove voxels of lesions outside the white and grey matter brain tissue, lesion volumes were filtered by means of custom masks based on the ICBM152 template.

### 5. Disconnectome Maps

Disconnectome maps were computed with the ‘disconnectome map’ tool of the BCBToolkit software (9). The first step of the procedure is the tracking of white matter fibres passing through each patient’s lesion, by means of the registration of lesions on the diffusion weighted imaging dataset of 10 healthy controls (39). This produces a percentage overlap map that takes into account the inter-individual variability of tractography in healthy controls’ dataset (40). Therefore, in the resulting disconnectome maps computed for each lesion, voxels show the probability of disconnection from 0 to 100% (11). These disconnection probabilities of each patient are then used for statistical analyses.

### 6. Statistical analysis producing the sites of lesion and tract disconnection

We ran 2 separate regression analyses for lesion sites and tract disconnections, using the same procedure. We used the tool “randomize” (41), part of FSL package (http://www.fmrib.ox.ac.uk/fsl/, version 5.0), which performs nonparametric statistics on neuroimaging data. Lesion drawings or disconnectome maps were taken into account as dependent variables within the general linear model implemented in ‘randomize’, in order to test the difference between the two groups in terms of disconnected brain regions. Demographic (age, education), clinical (lesion size, lesion onset- assessment interval, motor deficit) and neuropsychological (personal and extrapersonal neglect and memory impairment) data were considered in the model as covariates of non-interest. Threshold-Free Clusters Enhancement option was applied as to boost cluster-like structures of voxels and results that survived 5000 permutations testing were controlled for family-wise error rate (p>0.95).

### 7. Comparison of disconnection results with a brain atlas

In order to confirm the matching between white matter disconnection emerging from regression and the anatomy of each single tract, we used an atlas of human brain connections (42).

Maps of this atlas were first thresholded at 90% (i.e. the tract position in at least 90% of healthy population) and binarized to produce masks representing the cingulum, the frontal aslant (FAT) and the fronto-striatal tracts (FST) as well as the third branch of the superior longitudinal fasciculus (SFL III).

Then, these masks were used to extract the probabilities of disconnection for each tract from each patient’s disconnectome map. These probabilities were used to investigate the contribution of each tract and their disconnection co-occurrence.

### 8. Integration among tracts

To confirm the individual involvement of each tract and test their joint contribution to AHP, statistical analyses were conducted by using Bayesian models (R software, 43; brms package, 44) and generalised linear multilevel models were computed (Stan, 45).

The presence of AHP (1) or its absence (0) was used as the dependent variable, while keeping demographic, clinical and neuropsychological variables as covariates of non-interest. As covariates of interest, we used the probability of each tract disconnection, ranged between 0 (=no lesion) to 1 (=full lesion).

Then, we fitted 95 binomial models, starting from the null model (i.e., with only the covariates of non-interest) to the full model (i.e. with all the covariates of non-interest, the tracts, and all the interactions among them). The posterior samples were obtained by 4 chains, with 2500 burn-in and 2500 sampling iterations, resulting in a total of 10000 iterations for each posterior sample.

As a first step we tested whether the single tracts can explain the presence of AHP better than the null model. For this, we used the Bayes Factor (BF_10_, 17). A BF_10_ greater than 3 shows positive support for the hypothesis that the tract is necessary, a BF_10_ greater than 150 shows very strong support (17).

Then, all the models were compared among them by means of their marginal likelihood, showing which is the winning model with a probability from 0 (=no representative) to 1 (=the best model).

### 9. Data availability

The raw data used for this research (lesions) as well as the dependent variable and covariates are provided in full as supplementary data.

